# Exploring Healthy Neurocognitive Ageing with Deep Learning Interpretability Methods

**DOI:** 10.64898/2026.09.14.751405

**Authors:** Quentin Sénant, Clément Guichet, Nicolas Grivel, Monica Baciu, Martial Mermillod

**Affiliations:** Univ. Grenoble Alpes, CNRS LPNC UMR 5105, Grenoble, 38000, France; Neurology Department, CMRR, Grenoble Hospital, Grenoble, 38000, France

**Author notes:** These authors contributed equally to this work (Quentin Sénant, Clément Guichet).

**Keywords:** Healthy ageing, Functional Connectome, Brain Functional Networks, Lifespan, Brain Age Prediction, Multilayer Perceptron, Attribution

## Abstract

Healthy ageing reorganises large-scale functional brain networks. Using resting-state fMRI from 615 adults aged 18 to 88 years in Cam-CAN. We combined raw functional-connectivity analyses, graph-theoretical topology, Ridge and MLP age prediction and their attributions in order to understand how functional brain organisation varies with age and relates to cognitive performance. Both models predicted age accurately, with a modest advantage for the MLP. Within-network FC decreased with age, while between-network FC and participation increased. Topological analyses showed increased participation and a redistribution of nodes across connector, provincial, satellite, and peripheral roles. The MLP SmoothGrad attributions emphasised high-participation and weakly specialised nodes, whereas counterfactual tests showed that replacing CON-SMN, CON-DMN and DMN-FPN connections with a young-adult template produced the largest reductions in predicted age. Brain-cognition PLS linked these mechanisms to a broad cognitive-performance axis and a secondary semantic/connectomic axis. Together, these findings suggest that healthy ageing involves a redistribution of functional organisation that extends beyond a single DMN-FPN axis towards a broader DMN-CON-SMN-FPN configuration.

## 1 Introduction

Healthy ageing reshapes large-scale brain organisation. Resting-state functional connectivity (FC) studies, including recent empirical and review work (Abellaneda-Pérez et al., 2019; Deery et al., 2023), show that within-network connectivity often decreases with age, whereas between-network coupling tends to increase. Researchers disagree on whether this dedifferentiation mainly reflects decline or compensation. Reduced specialisation may index lower neural selectivity and functional efficiency (Goh et al., 2010; Malagurski et al., 2020; Park et al., 2004; Stumme et al., 2020), whereas increased between-network integration may support compensatory recruitment across distributed systems (Cabeza et al., 2018; Park & Reuter-Lorenz, 2009; Reuter-Lorenz & Park, 2024). This may allow older adults to maintain cognitive performance, particularly in tasks with significant cognitive control demands (Cabeza et al., 2018; Park & Reuter-Lorenz, 2009; Reuter-Lorenz & Park, 2024).

Current evidence suggests that maladaptive and compensatory functional trajectories can develop simultaneously within the same individual (Cassady et al., 2020; Chen et al., 2022; Deschwanden et al., 2025; Lemoine, 2020; McDonough et al., 2022). For example, the Default-Executive Coupling Hypothesis of ageing (DECHA) suggests that a more dedifferentiated recruitment of the Default Mode Network (DMN) in older adults leads to a more rigid coupling with the brain’s executive control system, the Fronto-Parietal Network (FPN). This DMN-FPN rigidity may yield divergent cognitive outcomes depending on task demands: it promotes cognitive resilience in tasks that require prior semantic knowledge, but drives cognitive decline in tasks with high control demands. This fits with the broader view that the healthy ageing brain shifts towards a more “semanticised” form of cognition that operates with reduced control demands (Koshino et al., 2023; Spreng & Turner, 2019; Spreng et al., 2018; Turner & Spreng, 2015). Recently, the SENECA model has extended this perspective, suggesting that this shift towards more semantic processing in ageing is supported by perceptuo-motor and control-related systems such as the sensorimotor network (SMN) and the cingulo-opercular network (CON), a system involved in salience detection and stable task-set maintenance (Dosenbach et al., 2008). Therefore, age-related reorganisation may involve a broader reconfiguration involving the CON and the lower-level SMN (Guichet, Banjac et al., 2024; Guichet et al., 2026a).

Overall, current theoretical frameworks provide a rich description of the large-scale brain network reorganisation and its impact on cognitive ageing. However, the field of neurocognitive ageing relies predominantly on classical FC analyses, leaving emerging machine and deep learning methods largely unexplored. Conversely, deep learning has successfully localised relevant features in multiple neuroimaging modalities, such as structural MRI and FC, for classification or regression tasks (Böhle et al., 2019; Kim & Ye, 2020; Lei et al., 2022; McClure et al., 2023; Song et al., 2024; Sturm et al., 2016; Thomas et al., 2019; Zhang et al., 2021), their integration into theories of cognitive ageing has remained limited.

Consequently, it remains unclear whether advanced interpretability methods, derived from the deep learning literature, can reveal biologically plausible mechanisms that support or extend current ageing models.

Brain-age prediction models provide an entry point to bridge the gap between cognitive ageing theories and such interpretability methods: they predict chronological age from neuroimaging features and, when combined with attribution methods like SmoothGrad (Smilkov et al., 2017), can estimate how sensitive the model’s prediction is to individual input features (Rahman et al., 2022). This can therefore reveal connectomic patterns that carry age-relevant information, providing both advanced interpretability and biological relevance. Specifically, neuroscientific inferences may further gain in robustness if triangulated across multiple interpretability methods or systematically compared against alternative baseline controls.

### 1.1 Present study and hypotheses

Using resting-state fMRI data from the Cam-CAN cohort (Cambridge Centre for Ageing and Neuroscience; Cam-CAN et al., 2014; Taylor et al., 2017), we asked whether healthy ageing is associated with a multiscale redistribution of functional organisation, whether these effects are predictable by linear and nonlinear models, and whether model-derived age-relevant features map onto theoretically meaningful connectomic and cognitive dimensions. The dataset, preprocessing, and connectome construction are described in Section 2.1. We used age prediction as an entry point for interpretation, combining predictive modelling (Section 2.2), edge-, node-, and network-level connectome analyses (Section 2.3), and brain-cognition analyses based on cognitive PCA and PLS (Section 2.4).

Guided by SENECA and related models of cognitive ageing, we formulated four main hypotheses. (i) First, we expected ageing to be associated with reduced functional segregation and increased between-network coupling, observable in raw FC and graph-theoretical topology. (ii) Second, because these effects were expected to be broad and distributed across the connectome, we expected chronological age to be strongly predictable by a regularised linear Ridge model; the MLP was therefore expected to provide, at most, a modest predictive advantage, while offering a differentiable function for gradient-based and counterfactual interrogation. (iii) Third, we expected MLP attribution to emphasise integrative topological properties, reflected in positive associations with participation coefficient and negative associations with within-module *z*-score. We further tested whether these topology-attribution relationships varied with age. (iv) Fourth, we expected age-relevant connectomic mechanisms to extend beyond a single DMN-FPN axis towards a broader DMN-CON-SMN-FPN configuration, and to covary with latent cognitive axes, especially those reflecting broad cognitive performance and SENECA-relevant cognitive trajectories (i.e., declining cognitive control and increased semantic access with age).

## 2 Method

### 2.1 Participants, preprocessing, and connectome construction

We used resting-state fMRI data from 615 healthy adults from the Cambridge Centre for Ageing and Neuroscience cohort (Cam-CAN; Cam-CAN et al., 2014; Taylor et al., 2017). Participants ranged from 18 to 88 years of age (*m* = 53.9, *sd* = 18.4); the sample included 301 females and 314 males. Data were preprocessed using fMRIPrep (Esteban et al., 2019).

We used each participant’s T1-weighted image for skull stripping and registered the Schaefer-400 atlas (Schaefer et al., 2018) to native anatomical space with SyN registration in ANTs (Avants et al., 2011). We transformed atlas labels using nearest-neighbour interpolation. We then regressed nuisance signals using high-pass filtering, motion correction, white matter/CSF regression, global signal regression following Wang et al. (2024), and adaptive scrubbing. Finally, we extracted regional time series with Nilearn’s NiftiLabelsMasker (Abraham et al., 2014), using *z*-score standardisation and 6-mm FWHM spatial smoothing.

For each participant, we computed Pearson correlations between all regional time series. We vectorised the upper triangle of each 400×400 FC matrix, excluding the diagonal. This yielded 79,800 edge features per participant, which we used as model inputs.

We assigned Schaefer parcels to eight networks using the Ji et al. organisation (Ji et al., 2019): Auditory (AUD), Cingulo-Opercular (CON), Dorsal Attention (DAN), Default Mode (DMN), Frontoparietal (FPN), Language (LAN), Sensorimotor (SMN), and Visual (VIS).

We assigned participants to fixed age-stratified training, validation, and test sets of 429 (≈ 70%), 93 (≈ 15%), and 93 participants, respectively. We used the validation set for hyperparameter selection and early stopping, the test set for final hold-out evaluation, and 10-fold cross-validation to estimate predictive stability.

### 2.2 Predictive modelling framework

We trained an MLP and a Ridge regression on the same 79,800 FC features. Ridge provided a linear benchmark suited to high-dimensional, collinear connectomic data (Dadi et al., 2019; He et al., 2020; Zhou et al., 2023). The MLP modelled nonlinear relationships. We implemented it in PyTorch (v2.10.0; Paszke et al., 2019) with three fully connected layers and a linear age-prediction head.

We selected hyperparameters with Optuna and a Tree-structured Parzen Estimator sampler (Optuna v4.7.0; Akiba et al., 2019; Bergstra et al., 2011). The search used 80 trials, including 20 random startup trials, and minimised validation MAE. The final architecture used two 128-unit hidden layers, a 32-dimensional latent layer, ReLU activations, dropout of 0.04, batch size of 8, learning rate of 2.0 × 10*^−^*^4^, Gaussian input-noise standard deviation of 0.02, and an explicit L2 penalty *λ*_L2_ = 2.6 × 10*^−^*^4^.

We trained the MLP with AdamW (Loshchilov & Hutter, 2019) and set AdamW weight decay to zero, so that our explicit L2 term controlled shrinkage:

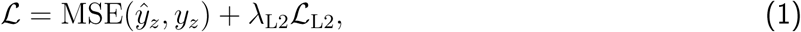

where *y_z_* and *y*^*_z_* denote observed and predicted standardised age. We standardised age within each training fold and converted predictions back to years. During training, we added Gaussian input noise and clipped noisy connectomes to the valid Pearson-correlation range [−1, 1]. We trained the final model on the fixed training set with early stopping and evaluated it on the independent test set.

The Ridge model was trained on the same input. Ridge regularisation strength was selected by minimising validation MAE across 17 logarithmically spaced values in the range [10*^−^*^3^, 10^5^], and by minimising validation MAE. In cross-validation, Ridge hyperparameter selection was repeated inside each fold. To compare MLP and Ridge performance across matched folds, we computed fold-wise differences so that positive values indicated better MLP performance:

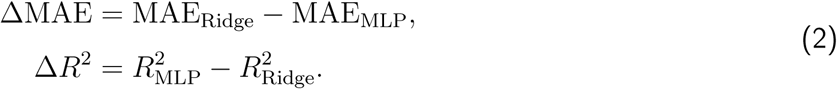

Model-derived edge importance was estimated using SmoothGrad (Smilkov et al., 2017). For each participant, we generated *K* = 50 noisy samples around the original input with a standard deviation of 0.15. The modified connectomes were clipped between −1 and 1 to ensure that all input edge values remained within their valid range. Then, for each noisy sample, we computed the absolute gradient of predicted age with respect to each input edge and averaged these gradients across noisy samples.

Finally, each participant-level attribution vector was min-max scaled to [0, 1]. These attribution scores quantified local model sensitivity to each FC edge. We computed Ridge edge contributions as |*x_i_β_i_*| and summarised them with the same edge-, node-, and network-level procedures.

### 2.3 Multilevel connectome analyses

We analysed the connectome at three levels. Edge-level analyses tested age-related FC changes. Node-level analyses characterised graph-theoretical roles and related them to model attribution. Network-level analyses summarised within-and between-system effects and tested their predictive contribution with ablation and counterfactual procedures.

#### Edge-level analyses

We first characterised age-related changes in raw FC. At the participant level, we computed global mean FC and tested its association with chronological age using Pearson and Spearman correlations. Statistical significance was assessed using age-label permutation across participants with 1,000 permutations.

At the edge level, we computed Spearman correlations between chronological age and each of the 79,800 FC edges. These edge-wise age effects were then summarised according to the large-scale network labels of the two incident parcels. For each network pair (*A, B*), we computed the mean age-connectivity association across all corresponding edges. Within-network effects corresponded to *A* = *B*, whereas between-network effects corresponded to *A* ≠ *B*. This allowed us to test whether age-related FC changes were preferentially expressed as reduced within-network segregation or increased between-network coupling.

To assess whether the observed network organisation of age-related edge effects exceeded what would be expected from spatially autocorrelated cortical organisation, we used spatial spin permutations. Parcel coordinates were obtained from the Schaefer-400 atlas using Nilearn. For each spin permutation, ROI network assignments were spatially rotated and reassigned. We then recomputed the within-network mean, between-network mean, within-minus-between contrast, and selected theory-driven network-pair summaries. Spin-based *p*-values were computed from 1,000 spatial permutations.

Raw FC, SmoothGrad attribution, and Ridge contributions were also summarised at the network-pair level. For each participant and each pair of networks, we computed the mean value of all edges linking the two systems. These summaries were used to visualise age-group differences, to compute local-versus-long-range transition indices, and to build the brain-side variables used in the PLS analyses.

#### Node-level analyses

Consistent with the procedure used in Guichet, Banjac et al. (2024), we characterised the role of each node from two graph-theoretical indices: the participation coefficient, indexing cross-module integration, and the within-module *z*-score, indexing within-module segregation. For each participant, the FC matrix was converted to absolute weights and thresholded using a proportional graph density of 10%. This threshold was used to remain consistent with previous work applying the same topological-role framework (Guichet, Banjac et al., 2024). To assess whether the conclusions depended on this choice, we repeated the topological-role analyses across proportional densities from 5% to 20% (Appendix A).

We then computed the participation coefficient and within-module *z*-score using the Python implementation of the Brain Connectivity Toolbox (BCTpy, v0.6.1; LaPlante, 2024; Rubinov & Sporns, 2010). Both metrics were standardised within participant, yielding *zPC* and *zWMZ*. Nodes were then assigned to one of four topological roles:

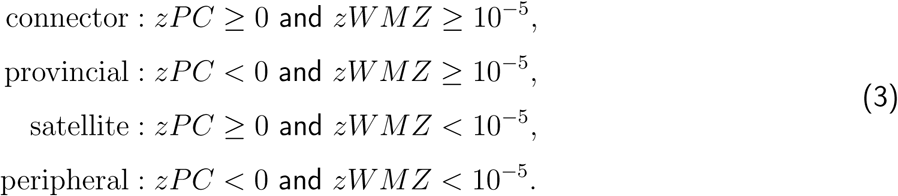

Thus, connector nodes combined high cross-module participation and high within-module strength, provincial nodes showed low cross-module participation but high within-module strength, satellite nodes showed high cross-module participation but low within-module strength, and peripheral nodes showed low values on both dimensions. We used these within-participant standardised values for topological-role assignment and the topology-attribution regressions, but retained the original continuous values for the age-trajectory analyses.

For each participant, we computed both the number and the proportion of connector, provincial, satellite, and peripheral nodes at the whole-brain level and within each resting-state network. Age trajectories of role proportions were analysed using quadratic binomial models with a logit link. For each role-by-network combination, the number of nodes assigned to the target role was modelled as a binomial count. The age trajectory was modelled as:

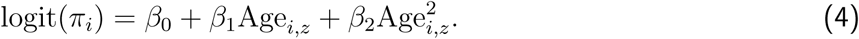

Alternative approaches include centred log-ratio transformations and multinomial models that treat the four role proportions jointly as compositional data. We used grouped quadratic binomial models because they provide directly interpretable role-specific trajectories while constraining fitted values to the unit interval. As a compositional control, we also fitted a grouped quadratic multinomial regression to the joint role counts. This control reproduced the main whole-brain and targeted-network trajectories. We fitted the binomial models separately for each role at the whole-brain level and within each large-scale network. Mean participation coefficient and mean within-module *z*-score were analysed separately using quadratic Gaussian models.

Node-level attribution was computed by averaging edge-level SmoothGrad scores across all edges incident to each node. Note that we did not use the degree formula here, as the SmoothGrad scores derived for each edge do not describe an adjacency matrix and thus do not permit the use of graph-theoretical tools. This quantified how strongly the model’s age prediction depended on each node’s connectivity profile, yielding one nodal attribution score per participant and parcel. The same edge-to-node aggregation was applied to Ridge contribution values. We then tested whether nodal importance was related to graph-theoretical integration and segregation by fitting the following linear model:

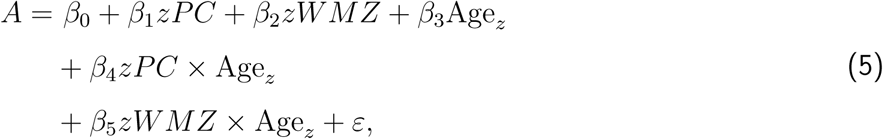

where *A* denotes nodal attribution or Ridge contribution. Age*_z_* denotes the standardised age of participant. Standard errors were clustered by participant to account for repeated nodal observations within individuals. We fitted the same model separately to absolute SmoothGrad attribution and Ridge nodal contribution values.

#### Network-level analyses

Network-level contributions to age prediction were assessed using three complementary approaches: chord-diagram visualisation, leave-one-network-out Ridge ablation, and targeted counterfactual interventions.

First, we visualised network-pair summaries using chord diagrams. We averaged the edge-wise Spearman age-FC correlation coefficients across all edges belonging to each network pair. For descriptive purposes, we also computed mean signed FC separately in young adults (<45 years, *N* = 212), middle-aged adults (45-65 years, *N* = 206), and older adults (>65 years, *N* = 197). These thresholds were chosen for two main reasons: they approximately divided the sample into age tertiles, thereby avoiding highly unbalanced groups; and the upper threshold placed the older-adult group above 65 years, a conventional chronological cut-off commonly used to define older or elderly adults (Kowal & Dowd, 2001). These age-group chord diagrams were used to visualise the average network-level organisation of the functional connectome across age groups.

Second, we performed a leave-one-network-out (denoted LO-RSN-O for leave one resting-state network out) Ridge ablation analysis. For each target resting-state network, all edges incident to that network were identified, including both within-network edges and between-network edges connecting the target system to any other system. These edges were replaced by their edge-wise mean values estimated from the training data, while all other edges were left unchanged. Ridge models were evaluated on original and ablated connectomes, and the loss of predictive performance was quantified using the reduction in *R*^2^ and the increase in MAE:

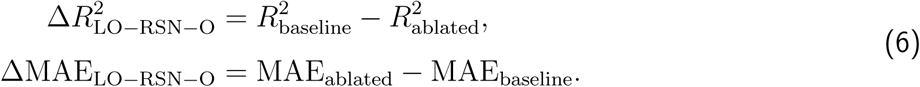

Because larger networks contain more incident edges, we normalised the LO-RSN-O effects per 1,000 replaced edges:

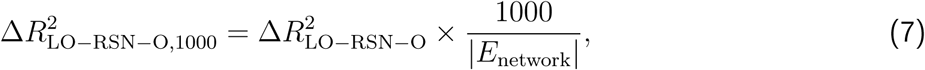

where |*E*_network_| is the number of replaced edges for the target network. The same normalisation was applied to MAE increases. We used 20 age-stratified cross-validation folds for this analysis in order to stabilise estimates. As an edge-count control, for each network and fold we generated an edge-count-matched null ablation by replacing the same number of randomly selected edges. Empirical LO-RSN-O effects were therefore interpreted relative to both edge-count-normalised values and edge-count-matched null effects.

Third, we performed targeted counterfactual age-prediction tests. To do so, we constructed a young-adult template by averaging the connectomes of training-set participants younger than 45 years. For each target network pair (*A, B*), we created a binary edge mask selecting either within-network edges when *A* = *B*, or between-network edges when *A* ≠ *B*. The target pairs focused on theoretically relevant interactions among the DMN, FPN, CON, and SMN, as identified by the SENECA model: DMN-FPN, DMN-DMN, FPN-FPN, DMN-SMN, CON-DMN, and CON-SMN.

For each participant and target network pair, we generated a counterfactual connectome by replacing only the selected edge set with the corresponding young-adult template values. The trained MLP was then applied to both the original and counterfactual connectomes. The counterfactual age-reduction effect was defined as

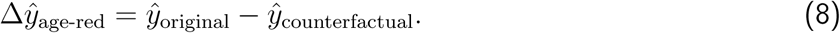

Positive values indicate that imposing young-adult connectivity values on the targeted edge set reduced predicted age. Counterfactual effects were also normalised per 1,000 replaced edges:

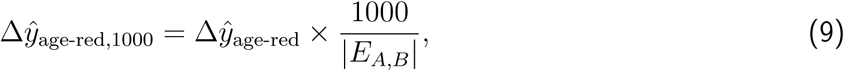

where |*E_A,B_*| is the number of replaced edges in the target network pair. The same counterfactual procedure was applied to the Ridge model as a linear control.

Finally, we computed a network-transition index designed to contrast a longer-range DMN-FPN component with a more local DMN-SMN-CON component. For each participant and each connectomic measure, we defined

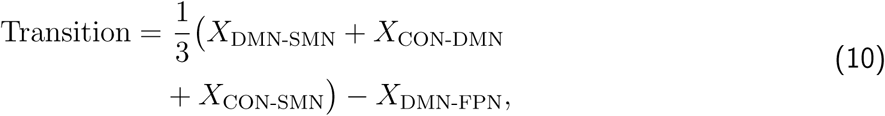

where *X* denoted raw FC, MLP SmoothGrad attribution, or Ridge absolute contribution.

### 2.4 Cognitive and brain-cognition analyses

We analysed eight neuropsychological measures available in Cam-CAN: Naming, Verbal Fluency, Proverb Comprehension, Sentence Comprehension, Tip-of-the-Tongue ratio, Hotel Task, Cattell fluid intelligence, and Story Recall. We imputed missing values using training-set means and standardised using training-set parameters. We fitted a principal component analysis (PCA) on the training set and applied the fitted transformation to all participants. The first two cognitive axes were retained. The first axis was oriented so that higher scores corresponded to better cognitive performance and lower scores to poorer cognitive performance. Age-related trajectories of cognitive axes were modelled using quadratic ordinary least-squares models. This PCA characterised the dominant latent dimensions independently of brain data.

To relate connectomic mechanisms to cognition, we performed a brain-cognition partial least-squares (PLS) analysis using the eight neuropsychological measures as separate cognitive-side variables. The PLS allowed us to identify cognitive dimensions that covary with connectomic organisation. This representation (i.e., using the eight cognitive scores instead of composite scores) preserved the specificity of each measure and allowed the cognitive dimensions to emerge without imposing theory-defined groupings. The brain-side PLS block included connectomic mechanisms derived from topological roles, raw FC, MLP SmoothGrad attribution, Ridge contribution, and counterfactual age-reduction effects. These variables focused on whole-brain participation and role proportions, DMN, FPN, CON and SMN participation and role measures, DMN-FPN, CON-DMN, CON-SMN, and DMN-SMN network-pair effects, and the local-minus-long transition indices.

We fitted the PLS models after mean imputation and standardisation of brain- and cognitive-side variables. For each latent variable, we reported the correlation between brain and cognitive PLS scores, the corresponding squared correlation, and permutation-based significance obtained by randomly permuting cognitive observations. Permutation testing used 2,000 permutations. Structure coefficients were computed as correlations between the original variables and their corresponding PLS scores, allowing interpretation of the variables contributing most strongly to each latent axis. Prediction was evaluated using 10-fold age-stratified cross-validation.

To clarify whether PLS2 represented a distinct brain-cognition axis, we performed three complementary checks focused on the raw FC variables included in the brain-side block. First, we fitted quadratic age models to each raw FC variable to quantify how strongly each network-pair measure tracked chronological age. Second, we inspected the structure coefficients of these raw FC variables on PLS1 and PLS2. Third, we tested whether each raw FC variable explained additional variance in brain and cognitive PLS scores after controlling for linear and quadratic age effects. This allowed us to determine whether PLS2 reflected the dominant age-related FC pattern.

Finally, we estimated descriptive path models linking age, brain PLS scores, and cognitive PLS scores for the first two latent variables. For each component, we fitted models with linear age alone and with linear and quadratic age terms. Indirect associations were estimated using 5,000 bootstrap samples. We reported percentile-based 95% bootstrap confidence intervals and two-sided bootstrap *p*-values.

### 2.5 Statistical inference and robustness checks

Statistical inference was adapted to the level of analysis. Subject-level associations were assessed with Pearson or Spearman correlations and, when appropriate, age-label permutation tests. Edge-level network-organisation effects were assessed using spatial spin permutations with 1,000 rotations to account for spatial autocorrelation of cortical parcels. Multiple comparisons across network-by-metric or role-by-network tests were controlled using false-discovery-rate correction. For binomial role-proportion models, model fit was summarised using deviance explained, denoted *D*^2^, which is a pseudo-*R*^2^.

For predictive modelling, the fixed hold-out test set was used only for final evaluation, while validation data were used for model selection and early stopping. Ten-fold cross-validation was used to estimate the stability of MLP and Ridge predictive performance. Paired MLP-Ridge differences were computed within matched folds, and confidence intervals were estimated across the 10 folds using the Student-*t* distribution.

For node-level topology-to-attribution models, standard errors were clustered by participant because each participant contributed repeated node-level observations. For LO-RSN-O and counterfactual analyses, both raw effects and effects normalised per 1,000 replaced edges were reported to avoid interpreting larger edge sets as intrinsically more informative simply because they contained more connections. LO-RSN-O effects were additionally compared with edge-count-matched random null ablations.

Finally, because graph-theoretical metrics can depend on proportional graph-density thresholds, we performed a BCTpy threshold-sensitivity analysis. Participation coefficient, within-module *z*-score, and nodal roles were recomputed across proportional densities of 5%, 7.5%, 10%, 12.5%, 15%, and 20%. For each threshold, the same quadratic binomial models were refitted for whole-brain and network-specific role proportions, whereas quadratic Gaussian models were refitted for continuous graph-theoretical metrics. Robustness was summarised using sign consistency relative to the primary 10% threshold, coefficient ranges across thresholds, stability of explained variance, and the number of thresholds for which effects remained significant after FDR correction.

## 3 Results

### 3.1 Predictive modelling framework

The regularised MLP generalised well to unseen participants, with *R*^2^ = .81, MAE = 6.57 years, and RMSE = 8.29 years on the independent test set. Ridge reached *R*^2^ = .78, MAE = 7.13 years, and RMSE = 8.82 years on the same participants (Table 1). Thus, the MLP reduced absolute prediction error but only modestly improved explained variance.

**Table 1:**
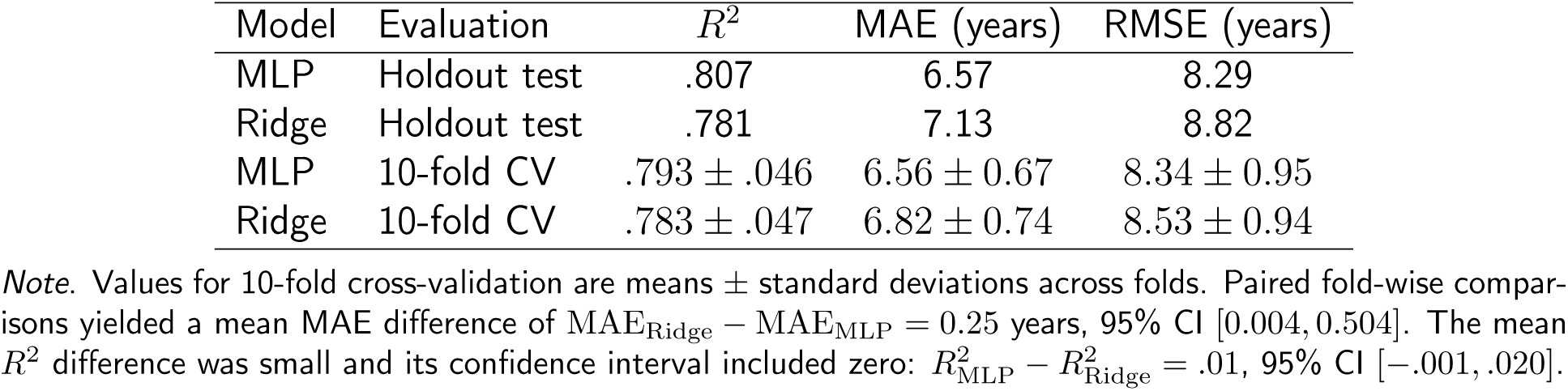
Age-prediction performance of the MLP and Ridge.

Ten-fold cross-validation showed the same pattern. The MLP had a small MAE advantage over Ridge, whereas the *R*^2^ confidence interval included zero. These results indicate that whole-brain FC strongly predicted chronological age and that a regularised linear model captured much of this signal.

These performances are consistent with published brain-age benchmarks, although direct comparisons remain limited by differences in modality, age range, and evaluation protocol. More et al. reported within-dataset test MAE values between 4.73 and 8.38 years and cross-dataset test MAE values between 5.23 and 8.98 years across systematic workflows (More et al., 2023). Their best Cam-CAN structural MRI workflow reached lower error than the present rs-FC models, but relied on voxel-wise structural features. Conversely, rs-fMRI-based studies summarised by Azzam et al. (2025) reported MAE values of 7.73, 5.14, and 8.24 years across different models and datasets. The present MLP and Ridge models therefore fall within the expected range for brain-age prediction from functional data.

### 3.2 Multilevel connectome analyses

#### Edge-level functional-connectivity ageing effects

In line with our first hypothesis, age-related FC effects reflected reduced functional segregation and increased between-network coupling. Mean whole-brain FC increased with age (Pearson *r* = .50, Spearman *ρ* = .53, both *p < .*001) and age-label permutation confirmed this association (*p*_greater_ = .001).

Edge-wise age analyses showed that age-related FC changes were organised by network structure. Across all edges, within-network connections showed a negative mean age correlation (*ρ* = −.05), whereas between-network connections showed a positive mean age correlation (*ρ* = .07). The resulting within-minus-between contrast was negative (Δ*ρ* = −.12) and significant under spatial spin permutation (*p*_spin_ = .001). This pattern supports an age-related reduction of within-network segregation together with increased between-network coupling. Thus, although the global mean FC increased with age, the edge-level decomposition showed that this global effect was primarily driven by increased between-network coupling rather than being uniform across all connections.

Targeted network-pair summaries showed the same organisation. Within-DMN and within-FPN edges showed negative mean age correlations (*ρ* = −.05 and *ρ* = −.05, respectively). In contrast, DMN-SMN, CON-DMN, and CON-SMN edges showed positive mean age correlations (*ρ* = .096, *ρ* = .092, and *ρ* = .085, respectively). The DMN-FPN mean edge-wise age correlation was close to zero (*ρ* = .004), indicating that age-related changes in the default-control axis were not expressed as a simple monotonic increase in raw DMN-FPN connectivity (Figure 2).

**Figure 1:**
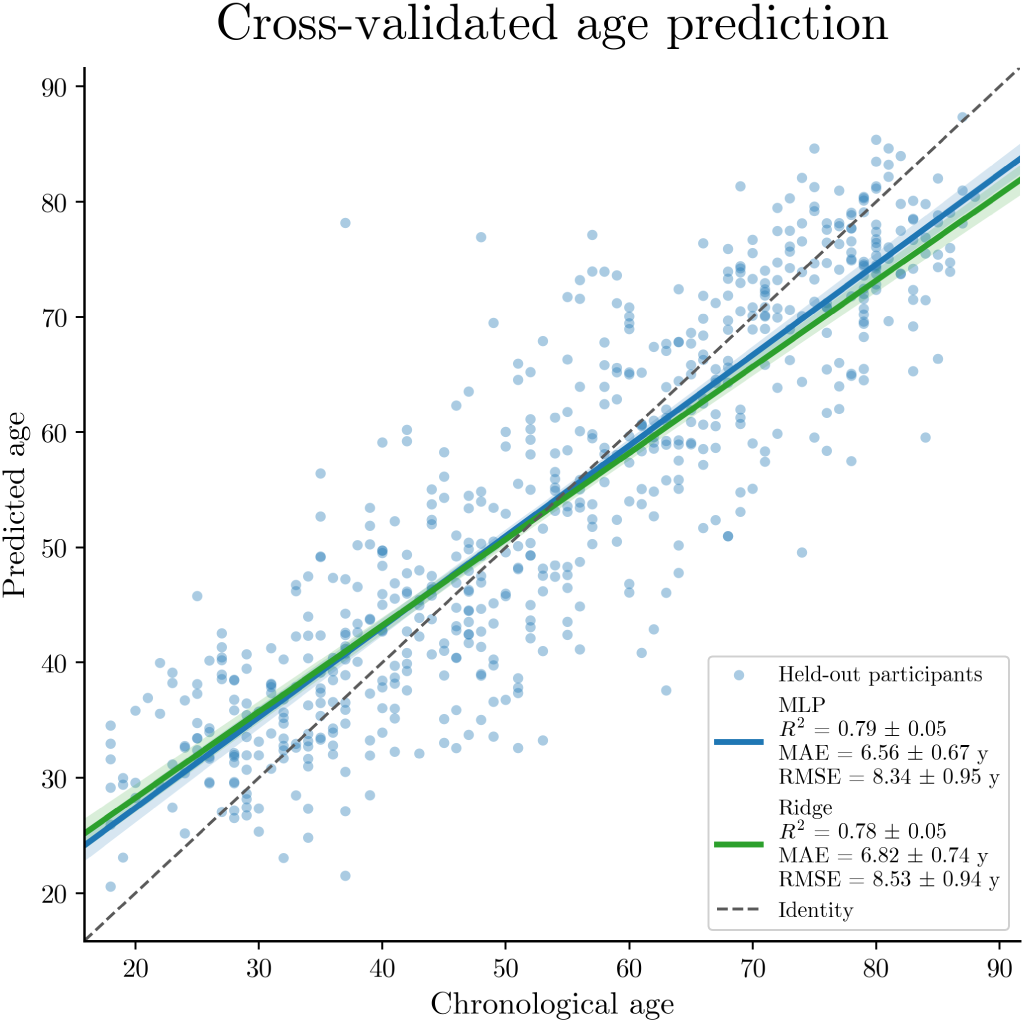
Chronological age versus out-of-fold predicted age in 10-fold cross-validation. Each point corresponds to a participant predicted only when held out from model training. The blue line and confidence band show the fitted MLP association; the green line and confidence band show the corresponding Ridge association. The dashed line indicates identity. Cross-validated *R*^2^, MAE, and RMSE are reported in the legend for each model.

**Figure 2:**
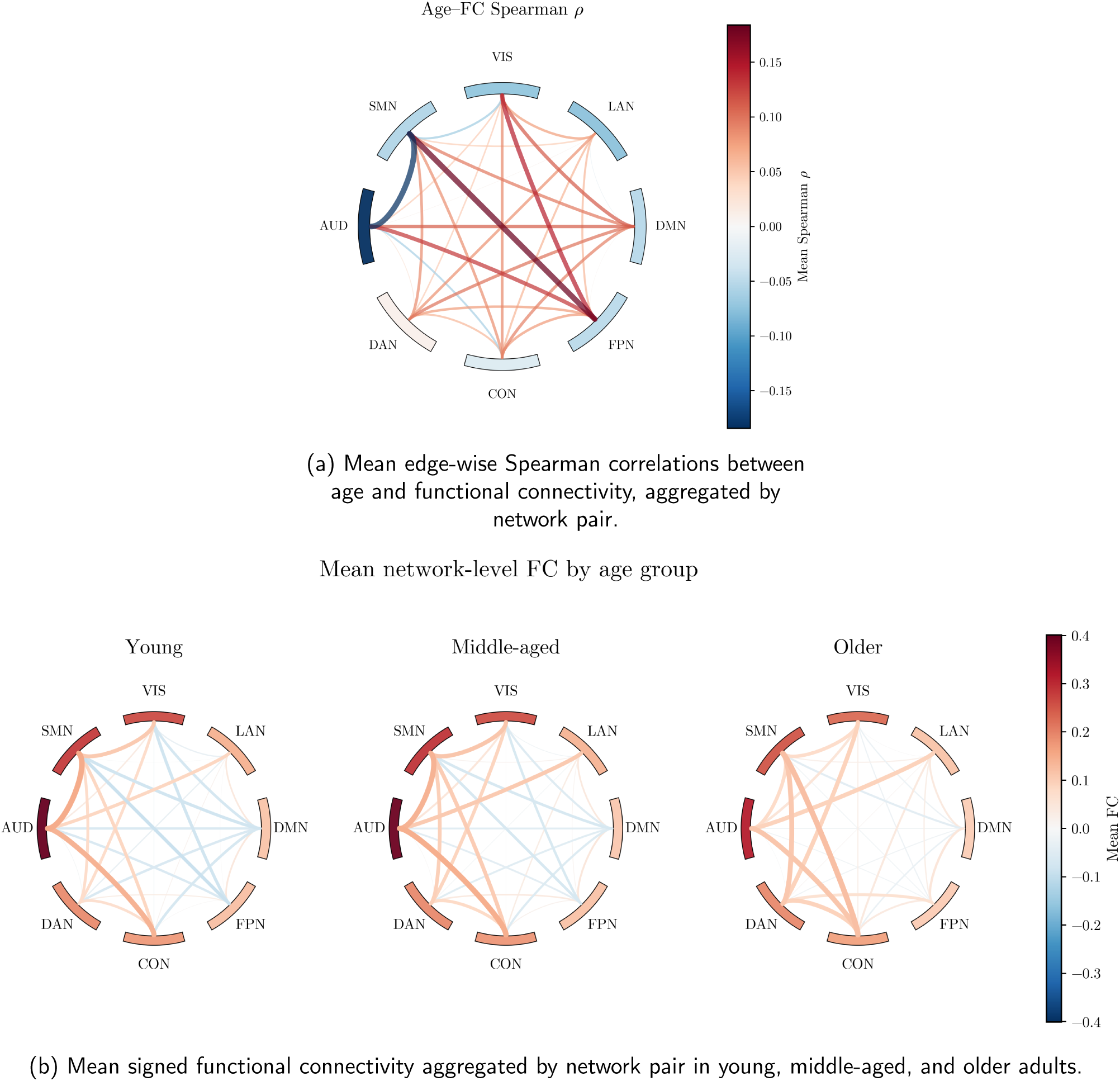
Network-level summaries of resting-state functional-connectivity ageing effects. *Note*. (A) Chord diagram summarising the primary edge-level ageing analysis. Each chord represents the mean signed Spearman correlation (*ρ*) between age and functional connectivity across all ROI-to-ROI connections belonging to a given network pair. (B) Descriptive chord diagrams showing the mean signed functional connectivity within each age group after aggregation by network pair. Outer arcs represent within-network mean FC and internal chords represent between-network mean FC. Network acronyms: AUD = Auditory network; CON = Cingulo-Opercular network; DAN = Dorsal Attention network; DMN = Default Mode network; FPN = Frontoparietal network; LAN = Language network; SMN = Sensor-imotor network; VIS = Visual network.

#### Node-level topological reorganisation

Topological roles were analysed at both the whole-brain level and the subsystem level. At the whole-brain level, quadratic binomial models showed significant age-related changes in all four role proportions. Connector proportion increased with age (deviance explained *D*^2^ = .06, *p*_FDR_ *< .*001), as did peripheral proportion (*D*^2^ = .07, *p*_FDR_ *< .*001). Conversely, provincial and satellite proportions decreased with age (*D*^2^ = .07 and *D*^2^ = .04, respectively; both *p*_FDR_ *< .*001). Whole-brain mean participation coefficient also increased with age (*R*^2^ = .15, *p*_FDR_ *< .*001). Thus, the dominant system-level topological signal of ageing was increased cross-module participation together with a redistribution away from provincial and satellite configurations.

Analyses at the subsystem level revealed heterogeneous role trajectories across networks. The CON showed increased connector and peripheral proportions, reduced provincial and satellite proportions, and increased mean participation (*R*^2^ = .08). The SMN showed the strongest increase in mean participation among the theoretically relevant systems (*R*^2^ = .19). Connector and satellite proportions increased, whereas provincial proportion decreased. Peripheral proportion showed no robust linear age effect. The DMN showed increased mean participation (*R*^2^ = .04), decreased satellite proportion, and increased provincial and peripheral proportions, indicating that its reorganisation did not correspond to a uniform shift towards connector-like organisation. The FPN also showed a mixed pattern: mean participation increased weakly but nonlinearly (*R*^2^ = .04), peripheral and provincial proportions increased, satellite proportion decreased, and connector proportion decreased. Thus, ageing reorganised nodal roles across systems with network-specific trajectories (Figure 3).

**Figure 3:**
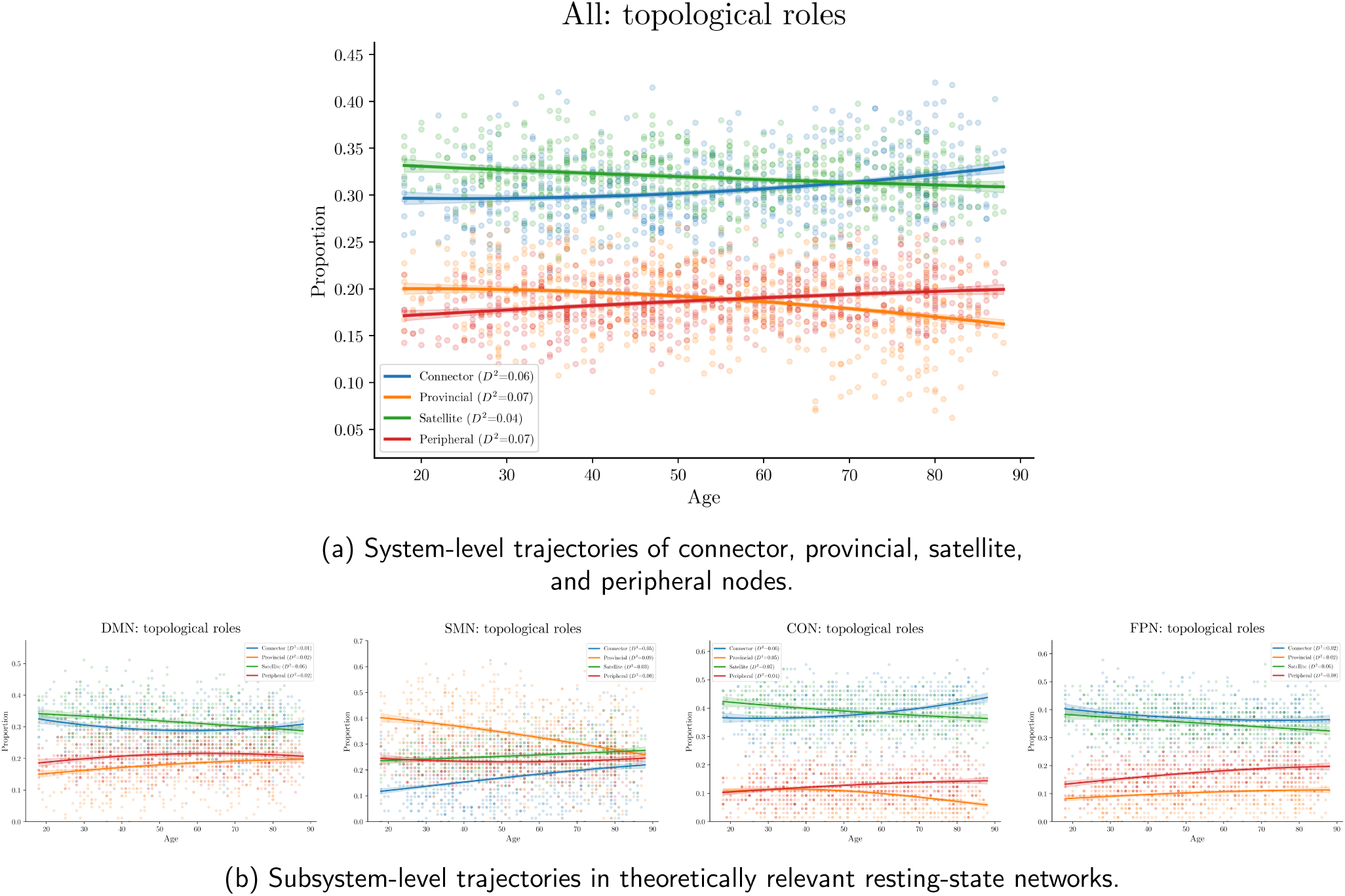
System-level and subsystem-level trajectories of topological roles across age. *Note.* Connector nodes combine high cross-module participation and high within-module strength; provincial nodes combine low participation and high within-module strength; satellite nodes combine high participation and low within-module strength; peripheral nodes show low values on both dimensions.

Threshold-sensitivity analyses confirmed that the main whole-brain topological findings were robust across proportional graph densities from 5% to 20%, whereas some network-specific role trajectories were more threshold-dependent (Appendix A).

#### Node-level attribution and graph topology

SmoothGrad maps were computed for all participants. Edge-level attribution scores were then averaged over all edges incident to each node, yielding one nodal attribution score per participant and parcel.

Absolute nodal SmoothGrad attribution was positively associated with participation coefficient (*b* = .02, *SE* = .003, *z* = 5.51, *p < .*001, 95% CI [.011*, .*023]). The MLP was therefore more sensitive to the connectivity profiles of nodes with greater cross-module participation. Absolute attribution was negatively associated with within-module *z*-score (*b* = −.16, *SE* = .002, *z* = −70.14, *p < .*001, 95% CI [−.167, −.158]). Thus, nodes with stronger within-module specialisation received lower absolute attribution.

Age was positively associated with nodal attribution (*b* = .28, *SE* = .006, *z* = 46.01, *p < .*001). The participation-by-age interaction was also positive (*b* = .01, *z* = 2.81, *p* = .005). The association between participation and attribution therefore became modestly stronger with age. The within-module *z*-score-by-age interaction was negative (*b* = −.01, *z* = −5.42, *p < .*001), indicating that the negative association between local specialisation and absolute attribution also strengthened with age.

The Ridge model showed a different topological profile. Ridge nodal contribution was positively associated with participation coefficient (*b* = .29, *z* = 58.85, *p < .*001) and within-module *z*-score (*b* = .38, *z* = 89.87, *p < .*001). The participation-by-age interaction was positive (*b* = .05, *z* = 9.76, *p < .*001). The within-module *z*-score-by-age interaction was small and did not reach significance (*b* = −.01, *z* = −1.90, *p* = .058).

Ridge contributions therefore reflected both integrative and locally specialised topology. In contrast, MLP SmoothGrad attribution preferentially highlighted high-participation and weakly specialised nodes. These complementary patterns suggest that the two models used different aspects of the same broad connectomic reorganisation.

#### Network-level contributions, transitions, and counterfactuals

The leave-one-network-out Ridge ablation tested how much age prediction deteriorated when all edges incident to a given network were replaced by their training-set mean values. In cross-validation, the largest per-edge *R*^2^ loss was observed for AUD 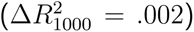, followed by CON 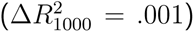, LAN 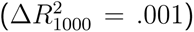, DMN 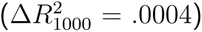, and SMN 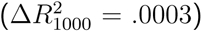. FPN and VIS showed smaller normalised effects, and DAN showed no evidence of a positive prediction loss. Thus, larger default-mode, control, and sensorimotor systems produced important absolute prediction losses, whereas the smaller auditory system showed the strongest per-edge effect.

The edge-count-matched null model confirmed that several empirical LO-RSN-O effects exceeded what would be expected from replacing the same number of arbitrary edges. Actual-minus-null *R*^2^ losses were largest for CON (.02), followed by DMN (.01), SMN (.007), LAN (.007), and AUD (.006). Together, the LO-RSN-O results indicate that age prediction relied on distributed network-level information, with particularly robust contributions from CON and DMN.

Network-pair transition analyses tested whether ageing was better described by a restricted DMN-FPN default-control axis or by a broader DMN-CON-SMN configuration. The local-minus-long transition index contrasted the mean of DMN-SMN, CON-DMN, and CON-SMN effects against the DMN-FPN effect. This transition explained *R*^2^ = .14 of the variance in raw FC, *R*^2^ = .69 in MLP SmoothGrad attribution, and *R*^2^ = .05 in Ridge absolute contribution.

For raw FC, the local-minus-long transition index increased linearly with age (*b* = .01, *t* = 10.00, *p < .*001), whereas the quadratic term was not significant (*b* = .002, *t* = 1.29, *p* = .196). For MLP SmoothGrad attribution, the same transition showed both a positive linear age effect (*b* = .002, *t* = 33.84, *p < .*001) and a negative quadratic term (*b* = −.001, *t* = −12.66, *p < .*001). Ridge absolute contribution showed a different and weaker profile, with a small negative linear effect (*b* = −2.3 × 10*^−^*^5^, *t* = −2.65, *p* = .006) and a negative quadratic effect (*b* = −4.2 × 10*^−^*^5^, *t* = −4.67, *p < .*001). Thus, as expected, MLP attribution emphasised a shift from DMN-FPN coupling towards broader DMN-CON-SMN organisation more strongly than Ridge contributions did (Figure 4).

**Figure 4:**
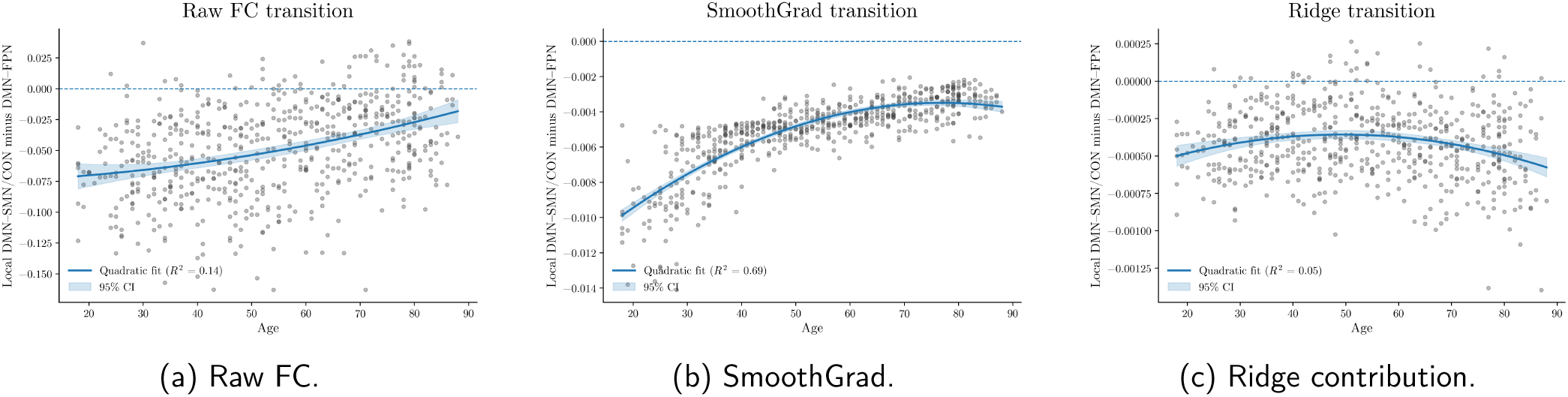
Age-related shift from DMN-FPN coupling towards broader DMN-CON-SMN organisation. Each panel shows the local-minus-long transition index contrasting local DMN-SMN/CON-DMN/CON-SMN interactions against the longer-range DMN-FPN component. Raw FC showed a linear age-related increase in this contrast, while MLP SmoothGrad attribution emphasised the same transition more strongly and nonlinearly. Ridge contribution showed a different and weaker pattern. Shaded bands indicate 95% confidence intervals.

Counterfactual analyses further supported a distributed network-level interpretation. Replacing targeted network-pair edges with a young-adult template reduced MLP-predicted age for all tested network pairs. After normalisation by the number of replaced edges, the largest MLP effects involved CON-SMN, CON-DMN, and DMN-FPN, with additional non-negligible effects for FPN-FPN and DMN-DMN. Ridge counterfactuals showed a related but not identical ordering, with the largest normalised effects involving CON-SMN, FPN-FPN, DMN-DMN, and DMN-FPN (Table 2).

**Table 2:** Counterfactual age-reduction effects normalised per 1,000 replaced edges.

| Network pair | Number of edges | MLP years / 1k edges | Ridge years / 1k edges |
| --- | --- | --- | --- |
| CON-SMN | 4680 | 0.241 | 0.297 |
| CON-DMN | 5590 | 0.237 | 0.259 |
| DMN-FPN | 5504 | 0.237 | 0.282 |
| FPN-FPN | 2016 | 0.203 | 0.287 |
| DMN-DMN | 3655 | 0.201 | 0.285 |
| DMN-SMN | 6192 | 0.096 | 0.100 |

*Note.* Values indicate the average reduction in predicted age per 1,000 replaced edges after replacing the targeted network-pair edges with the corresponding young-adult template values. Positive values indicate that the counterfactual connectome was predicted to be younger than the original connectome.

Counterfactual effects also increased with participant age. For the MLP, normalised age reductions were close to zero in young adults, increased in middle-aged adults, and were largest in older adults. This age-dependent pattern indicates that restoring young-like connectivity had the strongest model impact in participants whose original connectomes were most age-shifted (Figure 5).

**Figure 5:**
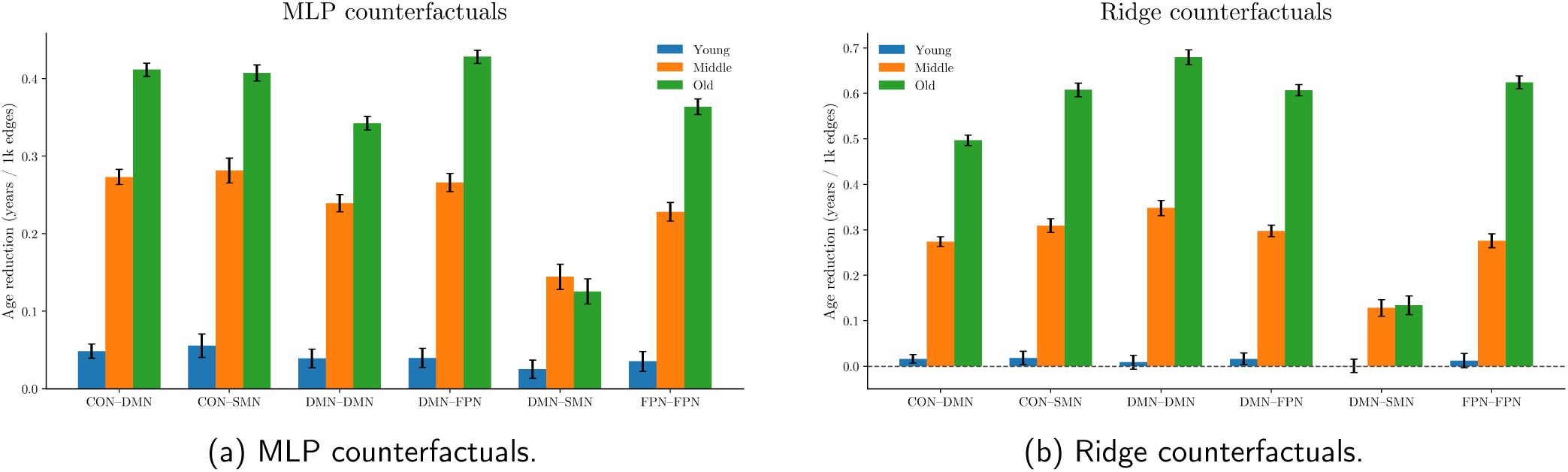
Counterfactual age-reduction effects normalised per 1,000 replaced edges. Positive values indicate that replacing the targeted network-pair edges with a young-adult template reduced predicted age. Error bars represent standard errors of the mean.

### 3.3 Cognitive and brain-cognition analyses

The first PCA-derived cognitive latent axis explained 35.4% of the variance across the eight neuropsychological measures. It loaded positively on all measures, with the strongest loadings for Naming (.43), Verbal Fluency (.42), Tip-of-the-Tongue ratio (.37), Story Recall (.37), Sentence Comprehension (.35), Cattell fluid intelligence (.31), Hotel Task (.29), and Proverb Comprehension (.25). It can therefore be interpreted as a broad cognitive-performance axis. A quadratic age model explained *R*^2^ = .39 of this axis, with both the linear age term (*b* = −.94, *p < .*001) and the quadratic age term (*b* = −.48, *p < .*001) significant. Thus, the dominant cognitive axis captured broad cognitive performance decline across adulthood.

The second cognitive axis explained 15.2% of the variance and showed a more semantic profile. It loaded positively on Proverb Comprehension (.67), Sentence Comprehension (.42), and Verbal Fluency (.15), and negatively on Cattell fluid intelligence (-.49), Naming (-.23), Tip-of-the-Tongue ratio (-.15), Story Recall (-.15), and Hotel Task (-.05). Therefore, it captured a more differentiated semantic dimension contrasting semantic comprehension with fluid and episodic-memory-related performance.

The brain-cognition PLS used the eight neuropsychological measures as separate cognitive-side variables and retained the first two latent variables (denoted PLS1 and PLS2). PLS1 showed a strong brain-cognition association (*r* = .66, *R*^2^ = .44, *p < .*001). On the brain side, this axis was mainly defined by MLP SmoothGrad attribution in DMN-SMN, CON-DMN, CON-SMN, and DMN-FPN, together with Ridge counterfactual effects in DMN-FPN, CON-DMN, and CON-SMN. On the cognitive side, it was mainly related to Naming, Cattell fluid intelligence, Story Recall, Tip-of-the-Tongue ratio, and Verbal Fluency. The structure coefficients showed that stronger expression of this connectomic axis covaried with lower broad cognitive performance. Cross-validated prediction was modest but non-negligible across the eight cognitive variables (*r* = .38, *R*^2^ = .140), with the strongest prediction for Cattell fluid intelligence, Naming, Story Recall, and Tip-of-the-Tongue ratio.

PLS2 also reached significance (*r* = .28, *R*^2^ = .08, *p < .*001). It was mainly defined by MLP and Ridge DMN-SMN counterfactual effects, raw DMN-SMN connectivity, and the local-minus-long FC index. On the cognitive side, it was dominated by Proverb Comprehension, Sentence Comprehension, and Verbal Fluency. PLS2 therefore captured a secondary semantic axis linked to DMN-SMN and local-minus-long connectomic variation.

The raw FC measures most strongly related to age were CON-DMN (*R*^2^ = .20), DMN-SMN (*R*^2^ = .16), the local-minus-long FC index (*R*^2^ = .14), and CON-SMN (*R*^2^ = .11), whereas DMN-FPN showed a much weaker age association (*R*^2^ = .01). PLS1 was aligned with this broad age-related FC pattern. In contrast, PLS2 was more specifically centred on DMN-SMN and local-minus-long variation (Figure 6). Finally, we fitted descriptive path models linking age, brain PLS scores, and cognitive PLS scores (Figure 7).

**Figure 6:**
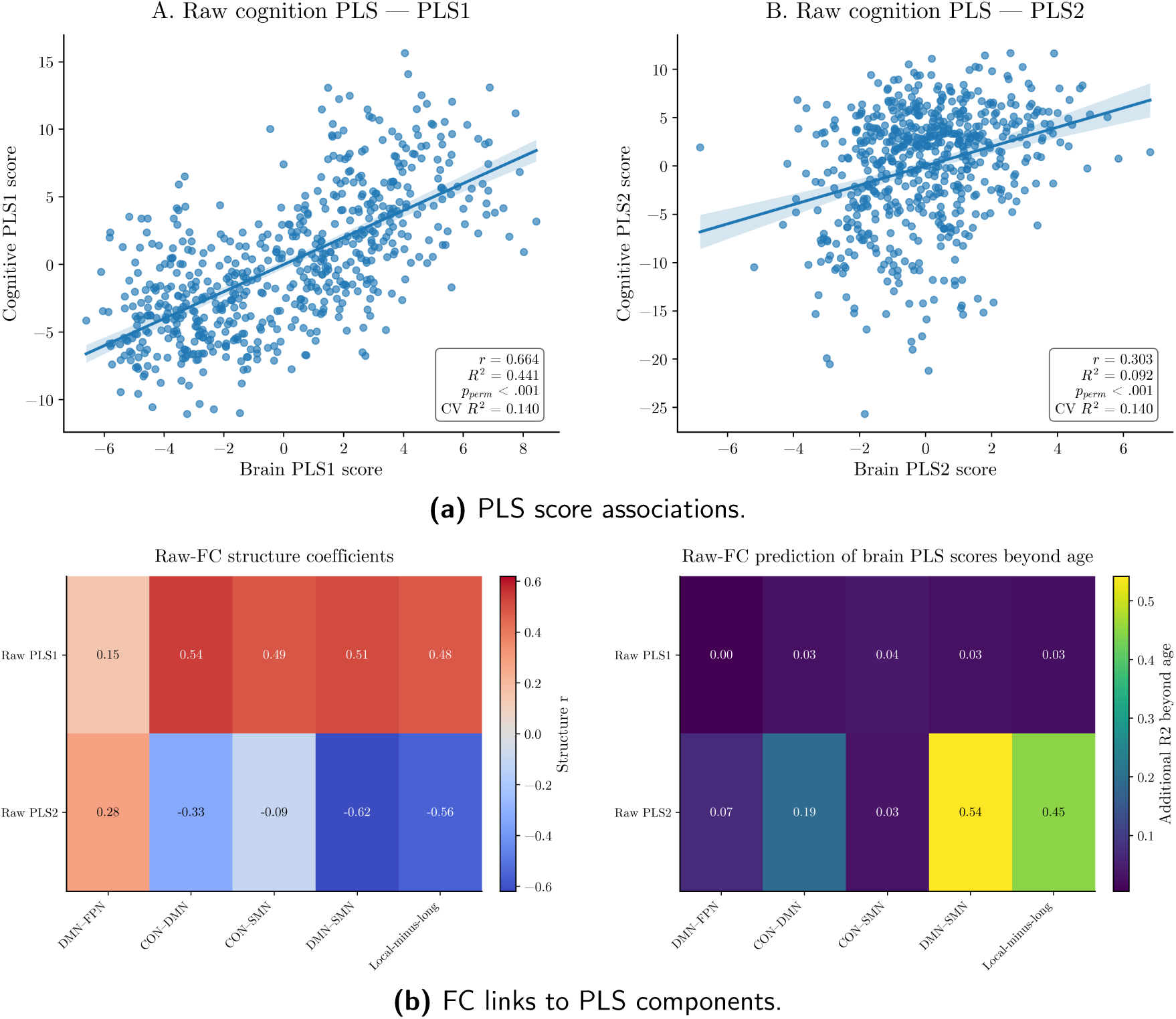
Brain-cognition PLS score associations and their relation to age-related raw functional-connectivity changes. **(a)** Associations between brain and cognitive scores for PLS1 and PLS2. The in-panel boxes report the score correlation, squared correlation, permutation-based significance, and cross-validated prediction performance. **(b)** Links between PLS components and age-related raw FC changes. The left heatmap shows the structure coefficients of raw FC variables on PLS1 and PLS2. The right heatmap shows the additional variance in brain PLS scores explained by each raw FC variable after controlling for linear and quadratic age effects.

**Figure 7:**
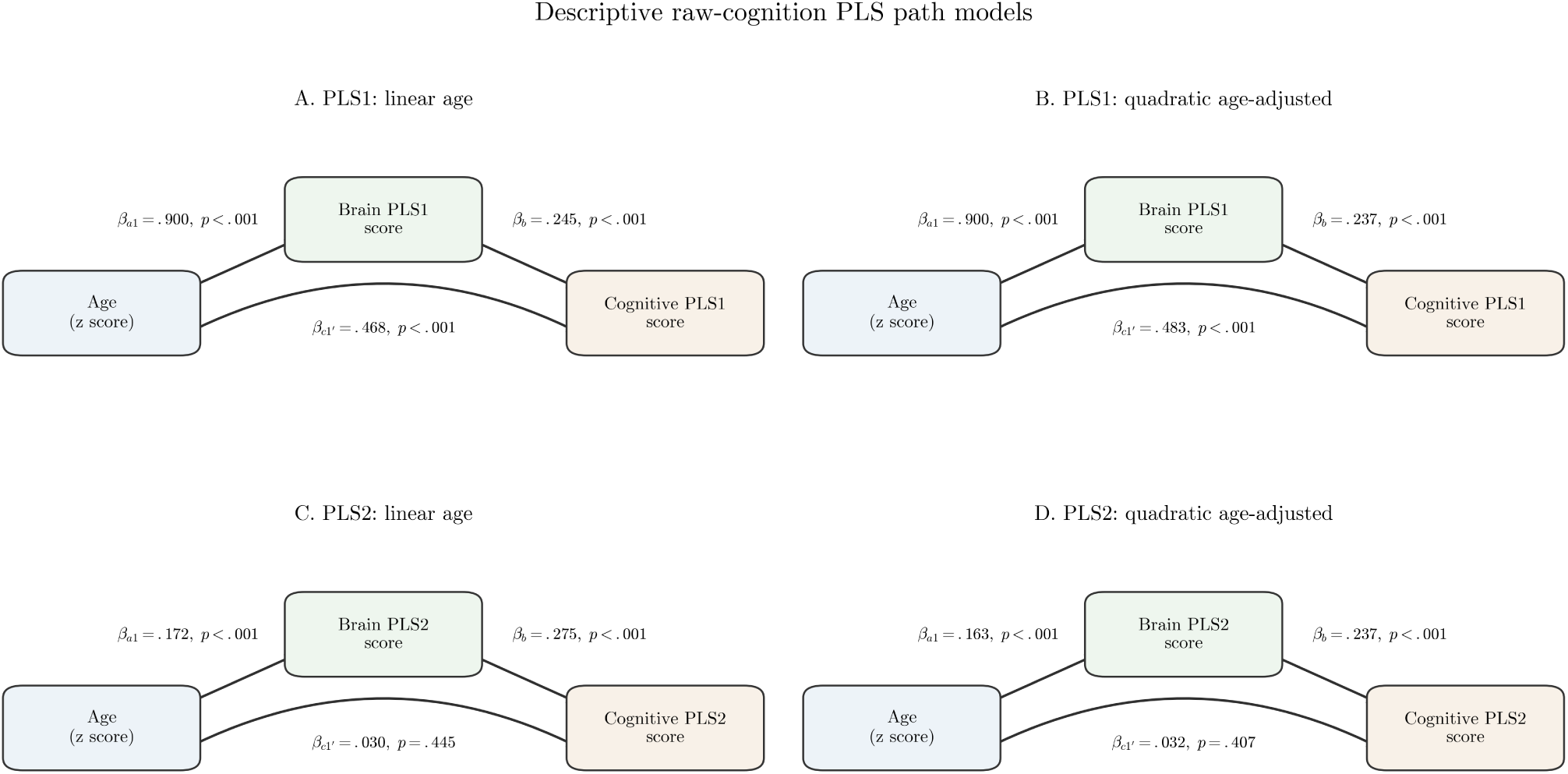
Descriptive age-brain-cognition path models for the first two PLS components. *Note.* Values shown beside the arrows are standardised path coefficients and their associated *p*-values. The path *a*_1_ represents the association between age and the brain PLS score, *b* represents the association between brain and cognitive PLS scores after adjustment for age, and *c^′^* represents the direct association between age and the cognitive PLS score after adjustment for the brain score. Linear and quadratic age terms were included in both models, although the quadratic paths are omitted from the diagrams for clarity.

In the quadratic age-adjusted PLS1 model, age strongly predicted the brain score (*β_a_*_1_ = .90, *p < .*001). Brain PLS1 remained associated with cognitive PLS1 after adjustment for linear and quadratic age (*β_b_* = .24, *p < .*001). Age also retained a direct linear association with cognitive PLS1 (*β_c_*_1_*′* = .48, *p < .*001). The linear indirect association was significant (*a*_1_*b* = .21, 95% CI [.11*, .*32], *p < .*001). Quadratic age showed little association with brain PLS1 (*β_a_*_2_ = .01, *p* = .722), but remained directly associated with cognitive PLS1 (*β_c_*_2_*′* = .27, *p < .*001). We found no evidence for a quadratic indirect association (*a*_2_*b* = .002, 95% CI [−.008*, .*011], *p* = .735). The model explained 81.0% of the variance in brain PLS1 and 54.7% of the variance in cognitive PLS1.

In the quadratic age-adjusted PLS2 model, age predicted the brain score (*β_a_*_1_ = .16, *p < .*001), and brain PLS2 remained associated with cognitive PLS2 (*β_b_* = .24, *p < .*001). The direct linear association between age and cognitive PLS2 was small and non-significant (*β_c_*_1_*′* = .03, *p* = .407). Nevertheless, the linear indirect association was significant (*a*_1_*b* = .04, 95% CI [.02*, .*06], *p < .*001). Quadratic age was negatively associated with brain PLS2 (*β_a_*_2_ = −.29, *p < .*001) and cognitive PLS2 (*β_c_*_2_*′* = −.15, *p < .*001). The quadratic indirect association was also negative (*a*_2_*b* = −.07, 95% CI [−.10, −.04], *p < .*001). The model explained 10.3% of the variance in brain PLS2 and 9.7% of the variance in cognitive PLS2.

In summary, the convergence of the cognition PCA and brain-cognition PLS on a broad-performance and a secondary semantic axis supported the interpretation of the PLS components that covary with age. The broad-performance axis was linked to distributed DMN-CON-SMN-FPN reorganisation and appeared predominantly linear with age, whereas the semantic axis was more specifically related to DMN-SMN and local-minus-long connectivity and showed a more pronounced quadratic trajectory.

## 4 Discussion

In the present study we used predictive modelling and attribution to test whether healthy ageing reorganises the functional connectome in ways predicted by current theoretical models of neurocognitive ageing. To do so, we examined four hypotheses. First, we expected ageing to reduce functional segregation and increase between-network coupling (i.e., functional dedifferenciation). Second, we expected nonlinear modelling to provide a small predictive advantage over linear modelling. Third, we expected MLP attribution to emphasise integrative rather than locally specialised topological properties. Fourth, we expected age-relevant connectomic mechanisms to extend beyond a single DMN-FPN axis towards a broader DMN-CON-SMN-FPN configuration and to covary with latent cognitive dimensions.

Across complementary analyses, the findings converged on an age-related reorganisation of the functional connectome. In line with the first hypothesis, ageing reduced within-network segregation, increased between-network coupling and participation, and redistributed nodal roles. In line with the second hypothesis, both Ridge and MLP predicted chronological age accurately, but the latter provided only a modest advantage, indicating that a regularised linear model captured much of the age-related connectomic signal. In line with the third hypothesis, SmoothGrad attribution emphasised high-participation and weakly specialised nodes, whereas contributions from the additive Ridge model reflected both integrative and locally specialised topology. Finally, in line with the fourth hypothesis, network ablations, counterfactual tests, transition indices, and PLS analyses converged on a distributed configuration involving the DMN, CON, SMN, and FPN. Taken together, these findings support current models of healthy ageing. We discuss the implications of this reorganisation below.

### 4.1 Multiscale connectome reorganisation

#### 4.1.1 System-level reorganisation

Age-related effects in raw FC followed the network structure. Within-network edges decreased with age, whereas between-network edges increased. This pattern supports the view that healthy ageing reduces functional segregation between specialised systems. Because this effect appeared before any predictive modelling, it provides a connectomic basis for interpreting the attribution and counterfactual analyses. Moreover, spatial spin testing confirmed that this organisation was significant.

This system-level profile suggests that ageing modifies the balance between local specialisation and cross-network integration. The concurrent increase in connector and peripheral configurations points to a heterogeneous redistribution of nodal roles rather than a uniform shift towards greater integration: some nodes participate more broadly across modules, whereas others become less strongly embedded within any single module. This pattern may emerge as locally specialised coupling weakens and coordination becomes distributed across a wider set of systems. The distributed neural efficiency account suggests that neural efficiency may depend on the selective allocation of processing resources across networks, producing local cost-efficiency trade-offs (Ramchandran et al., 2019). Older adults nevertheless tend to show lower segregation and modularity alongside greater integration, while also exhibiting lower local and global efficiency and weaker hub organisation (Deery et al., 2023). Increased cross-network integration should therefore not be equated with greater network efficiency. The simultaneous rise in connector and peripheral configurations may reflect a cost-constrained reorganisation in which broader cross-network coordination partly offsets reduced local specialisation, while producing a more diffuse and potentially less efficient architecture. This interpretation remains speculative. However, it is consistent with previous work suggesting that healthy ageing shifts functional organisation from locally specialised configurations towards more globally integrated, but still economically constrained, network states (Guichet, Banjac et al., 2024; Guichet et al., 2026a). The present findings cannot determine whether this redistribution primarily reflects compensation, dedifferentiation, or their coexistence.

The topological-role and attribution analyses extended this interpretation by linking age prediction to the balance between functional integration and specialisation. At the system level, connector and peripheral proportions increased with age, whereas provincial and satellite proportions decreased. Whole-brain participation coefficient also increased with age. These findings suggest that the ageing connectome undergoes a redistribution of nodal roles, with some nodes becoming more cross-module and others becoming less embedded in local modules. SmoothGrad attributions also showed that the MLP was especially sensitive to high-participation and weakly specialised nodes, whereas locally specialised nodes received lower absolute attribution. Thus, ageing may involve a shift away from highly segregated, domain-specific processing towards more distributed and integrative forms of network coordination. Such a shift may reflect dedifferentiation when reduced local specialisation indexes declining neural selectivity, but it may also support compensation when increased cross-network participation allows additional systems to be recruited to maintain performance.

#### 4.1.2 Network-level reorganisation

At the subsystem level, however, the pattern was not uniform. CON and SMN showed the clearest increases in participation and connector-like organisation, suggesting a prominent role in age-related cross-network integration. DMN and FPN followed more mixed trajectories, combining increased participation with changes in provincial, peripheral, or satellite roles. Thus, ageing altered integration and specialisation differently across resting-state networks.

The LO-RSN-O Ridge ablation provided a linear-model perspective on network importance. CON, DMN, and SMN produced the largest raw performance losses, whereas AUD showed the strongest per-edge effect after edge-count normalisation. CON and DMN remained robust across both perspectives, indicating that control and default-mode systems carried age-predictive information.

We then related MLP SmoothGrad attribution to graph-theoretical topology measures. Absolute nodal attribution was positively associated with participation coefficient and negatively associated with within-module *z*-score. The MLP was therefore more sensitive to integrative and weakly specialised topology. Both associations varied with age. The positive association between participation and attribution strengthened modestly with age. The negative association between within-module *z*-score and attribution also became stronger. Thus, the MLP increasingly emphasised integrative connectivity profiles relative to locally specialised profiles across adulthood.

In our results, raw FC showed little evidence of a strong monotonic increase in DMN-FPN coupling, and FPN integration followed a heterogeneous rather than uniformly increasing trajectory. These effects may depend on the spatial scale and network definitions used to summarise the connectome. Previous work has shown that the posterior cingulate cortex, and the DMN more broadly, is functionally heterogeneous. Its dorsal and ventral subdivisions differ in their connectivity and functional profiles and contribute differently to cognitive control. Moreover, their interactions with other brain systems vary with attentional state and processing demands (Leech & Sharp, 2014; Leech et al., 2011). Aggregating these distinct anatomical and functional subdivisions within a single label may therefore combine connections with different or even opposing age trajectories. This loss of functional resolution could attenuate age-related effects when connectivity is averaged across the entire DMN. The weak monotonic effect observed at this scale therefore does not preclude more spatially specific changes in default-executive interactions.

Counterfactual analyses supported the same distributed interpretation. Replacing selected edges with a young-adult template reduced predicted age across all tested network pairs. The largest normalised MLP effects involved CON-SMN, CON-DMN, and DMN-FPN, with additional effects in FPN-FPN and DMN-DMN. Ridge counterfactuals highlighted a related but differently ordered set of pathways. Thus, the MLP and Ridge converged on default-mode, control, and sensorimotor systems, but they weighted specific interactions differently. This suggests partial convergence between nonlinear and linear models, but also indicates that MLP attribution and counterfactual sensitivity may capture a somewhat different organisation of age-relevant CON-SMN and CON-DMN interactions. DMN-SMN connectivity was clearly age-related in raw FC analyses and contributed to the secondary PLS semantic axis, but it produced weaker counterfactual age-reduction effects than CON-SMN. Thus, DMN-SMN may index a more cognitive or semantic-perceptuo-motor dimension of ageing, whereas CON-SMN may provide a more directly age-diagnostic pathway.

In its initial formulation, SENECA emphasised the synergistic balance between FPN integration and DMN regulation as a key mechanism of cognitive ageing (Guichet, Banjac et al., 2024). Its revision broadened this account by proposing that age-related reorganisation also involves interactions with perceptuo-motor and control-related systems, particularly the SMN and CON (Guichet et al., 2026a). The present findings support this broader formulation. Ageing was associated with reduced segregation, increased cross-network participation, and a redistribution of functional importance across large-scale systems. Moreover, CON-DMN and CON-SMN interactions emerged as prominent components of age-related functional reorganisation.

Because the CON has been associated with salience detection, task-set maintenance, and stable control across cognitive contexts (Dosenbach et al., 2008; Power et al., 2011), its age-related involvement may reflect a shift in how the ageing brain coordinates internal, executive, and sensorimotor information. Strong CON-DMN and CON-SMN counterfactual effects suggest that age prediction depends on how control systems interface with default-mode and sensorimotor pathways. Within the SENECA framework, this may indicate that the brain increasingly relies on more distributed and possibly more constrained communication routes to maintain functional organisation under increasing biological constraints.

The PLS analysis linked age-relevant connectomic mechanisms to cognition. PLS1 captured a broad brain-cognition axis related to global cognitive performance, whereas PLS2 linked DMN-SMN and local-minus-long variation to a more semantic profile. The convergence of the PCA and PLS on these two dimensions strengthened the interpretation of the PLS components and indicated that the identified connectomic and cognitive profiles covaried across the ageing process. This pattern is consistent with the idea that DMN-SMN coupling may contribute to semantic access and retrieval in ageing, especially when cognition becomes more reliant on embodied, perceptuo-motor, or action-related representations (Guichet, Banjac et al., 2024; Guichet, Roger et al., 2024; Guichet et al., 2026b).

Finally, the path models showed distinct age-related patterns across the two latent variables. For PLS1, the dominant association was linear: age was strongly associated with the brain score, which remained associated with broad cognitive performance after age adjustment. The linear indirect association was significant, whereas the quadratic indirect association was negligible. PLS2 showed a weaker linear age association but a marked quadratic component. Both its linear and quadratic indirect associations were significant, with the quadratic component showing the opposite sign. This suggests that the secondary semantic brain-cognition axis follows a more pronounced nonlinear age-related trajectory than the broad PLS1 axis.

### 4.2 Relation to previous models of healthy ageing

Our results align with several models of cognitive ageing without directly testing all of their assumptions. We did not assess hemispheric asymmetry as proposed by HAROLD, posterior-to-anterior redistribution as proposed by PASA, or the load-dependent recruitment predicted by CRUNCH (Cabeza, 2002; Davis et al., 2008; Reuter-Lorenz & Cappell, 2008; Schneider-Garces et al., 2010). These frameworks were developed primarily from task-evoked neuroimaging, whereas we wanted to focus on the resting-state functional organisation. Recent studies have also questioned strong compensatory interpretations of these activation patterns (Haitas et al., 2024; Jamadar, 2020; Knights et al., 2021; Morcom & Henson, 2018; Myrum, 2019; Zając-Lamparska et al., 2024). Reduced hemispheric asymmetry and increased prefrontal recruitment may reflect less specific or less differentiated neural activity rather than successful compensation (Knights et al., 2021; Morcom & Henson, 2018; Myrum, 2019). Several direct tests have also failed to confirm the load-dependent activation pattern predicted by CRUNCH across working-memory and semantic tasks (Haitas et al., 2024; Jamadar, 2020; Zając-Lamparska et al., 2024). These findings challenge the assumption that age-related overactivation or redistribution necessarily reflects effective compensation.

Our findings are more directly relevant to the DECHA and SENECA frameworks. The involvement of DMN-FPN interactions is consistent with the importance of default-executive coupling proposed by DECHA. However, the strongest converging evidence also involved CON-DMN and CON-SMN interactions. This pattern suggests that age-related functional reorganisation is not limited to the regulation of the DMN by frontoparietal control systems. Instead, it may also involve cingulo-opercular and sensorimotor systems that support salience detection, task-set maintenance, bodily/perceptuo-motor grounding, and bottom-up control. In this respect, the present findings support the broader logic of SENECA while extending it beyond a single DMN-FPN axis towards a distributed DMN-CON-SMN-FPN configuration.

The involvement of CON-DMN and CON-SMN pathways points to an age-relevant architecture that extends beyond an isolated DMN-FPN axis. This four-system configuration is consistent with the shifting-architecture account proposed by Spreng and Turner (2019). According to this framework, ageing shifts cognition away from fluid and exploratory processing towards greater reliance on crystallised knowledge, semantic representations, and autobiographical experience. Our findings suggest a possible connectomic substrate for this transition. The DMN may support internally oriented and semantic representations, while the FPN enables flexible top-down selection. The CON may stabilise task sets and update control in response to relevant signals, whereas the SMN may ground cognitive representations in action, bodily state, and perceptuo-motor constraints.

This account also fits the exploration-to-exploitation model of late-life cognition (Spreng & Turner, 2021). Younger adults may rely more on flexible and controlled search, whereas older adults may increasingly exploit accumulated knowledge and familiar schemas. Within this framework, DMN-FPN coupling may support the controlled selection of internally generated knowledge. CON-DMN and CON-SMN interactions may then help stabilise this knowledge and translate it into action-oriented control states. This interpretation agrees with evidence that cingulo-opercular and frontoparietal systems make distinct contributions to cognitive control (Cao & Cannon, 2021; Cocuzza et al., 2020; Crittenden et al., 2016). In older adults, cingulo-opercular connectivity relates to several executive functions, whereas frontoparietal connectivity appears more specifically associated with working memory (Hausman et al., 2022). Recent accounts further describe the CON as an action-mode network involved in arousal, task initiation, action planning, and feedback-based updating (Dosenbach et al., 2025).

The SMN effects may therefore reflect more than low-level motor processing. They may indicate greater reliance on perceptuo-motor anchoring, bodily feedback, and familiar action routines, consistent with the functional coupling between somato-cognitive action regions and the CON (Gordon et al., 2023), as well as with evidence for distinct decision, action, and feedback components within the CON (D’Andrea et al., 2023). Such a configuration may support performance when older adults draw on stable and semantically rich representations, but may become less efficient during rapid switching or abstract control. We suggest that the DMN-CON-SMN-FPN architecture may therefore link compensation and dedifferentiation by providing additional control and sensorimotor support while reducing the separation between specialised systems.

### 4.3 Methodological implications

Ridge captured much of the age-related FC signal. The MLP added a modest predictive gain and a differentiable model for attribution. SmoothGrad highlighted integrative, weakly specialised nodes and a strong age association in the transition index. Its main contribution here is therefore to characterise predictive sensitivity alongside a strong linear benchmark. Moreover, all of our analyses pointed to a broad reallocation of functional importance across large-scale systems. We argue that this triangulation provides stronger support than a conclusion based on predictive performance or attribution maps alone.

### 4.4 Limitations and future directions

Several limitations should be considered. First, our design captures differences between people. Longitudinal data will be needed to determine whether changes in the brain connectome reflect progressive within-person reorganisation, cohort-specific differences, or both.

Analytical choices and participant characteristics may also affect the results. These include the parcellation, preprocessing strategy, global signal regression, graph threshold, etc. Although motion correction, nuisance regression, and adaptive scrubbing were applied, framewise-displacement-related variance may persist in resting-state FC and could partly overlap with age-related effects. Similarly, sex-related differences in brain organisation may contribute to individual variability in FC and brain-age prediction.

Future work should also examine whether age-related topological-role trajectories differ between anterior and posterior subdivisions of the DMN. Our analyses treated the DMN as a single large-scale system, whereas anterior and posterior DMN regions may contribute differently to semantic, autobiographical, control-related, and internally oriented processes (Leech et al., 2011). More recent work has further distinguished three DMN subsystems and shown that they make complementary contributions to semantic cognition (Shao et al., 2024). Separating these subsystems could therefore clarify whether age-related increases in participation, peripheralisation, or connector-like organisation are driven by specific DMN components rather than by the DMN as a whole. Such analyses would be particularly relevant for testing whether the weak raw DMN-FPN effect observed here reflects limited default-executive reorganisation or a loss of functional specificity caused by averaging connections with heterogeneous, distinct or even opposing age trajectories.

The functional meaning of the reorganisation also remains open. Resting-state associations alone leave compensation, dedifferentiation, and their coexistence unresolved. Task-based and longitudinal studies could distinguish these possibilities and directly test the predictions of HAROLD, PASA, and CRUNCH. A clinical extension could examine the brain-age gap as a marker of deviation from normative ageing. However, this application would require evidence that individual brain-age gaps predict clinically meaningful cognitive or neurological outcomes (Azzam et al., 2025; Baecker et al., 2021; Cole et al., 2017), which require validation in additional cohorts.

### 4.5 Conclusion

By triangulating evidence across analyses, this study provides robust evidence that healthy ageing is associated with reduced functional segregation, greater participation, and a redistribution of nodal roles in the brain connectome. This distributed reorganisation is linked to cognitive performance and a secondary semantic dimension associated with DMN-SMN and local-to-long-range connectivity. The resulting broad DMN-CON-SMN-FPN configuration supports a more distributed account of healthy neurocognitive ageing than accounts centred primarily on DMN-FPN coordination.

## Data and Code Availability

The authors do not have permission to redistribute the Cam-CAN data. The dataset is available upon request through the Cambridge Centre for Ageing and Neuroscience. The analysis code will be made publicly available upon submission.

## Declaration of the use of AI

AI played no role in planning or carrying out the research. AI (or other tools) were used to check spelling and grammar and to reword our draft text.

### 4.5 Conclusion

#### Author Contributions

**Quentin Sénant**: Writing – original draft, Writing – review & editing, Visualization, Validation, Software, Methodology, Investigation, Formal analysis, Data curation, Conceptualization.

**Clément Guichet**: Writing – review & editing, Validation, Software, Methodology, Investigation, Formal analysis, Data curation, Conceptualization.

**Nicolas Grivel**: Writing – review & editing, Methodology, Conceptualization.

**Monica Baciu**: Writing – review & editing, Validation, Supervision, Project administration, Funding acquisition.

**Martial Mermillod**: Writing – review & editing, Validation, Supervision, Project administration, Funding acquisition.

## A Sensitivity of topological-role analyses to graph-density threshold

To assess whether the topological findings depended on the proportional graph-density threshold used before BCTpy computation, we repeated the full topological-role analysis across six thresholds: 5%, 7.5%, 10%, 12.5%, 15%, and 20%. For each threshold, we recomputed the participation coefficient, within-module *z*-score, and the four nodal roles. We then refitted the same quadratic binomial models for role proportions and the same quadratic Gaussian models for continuous graph-theoretical metrics.

**Figure 8:**
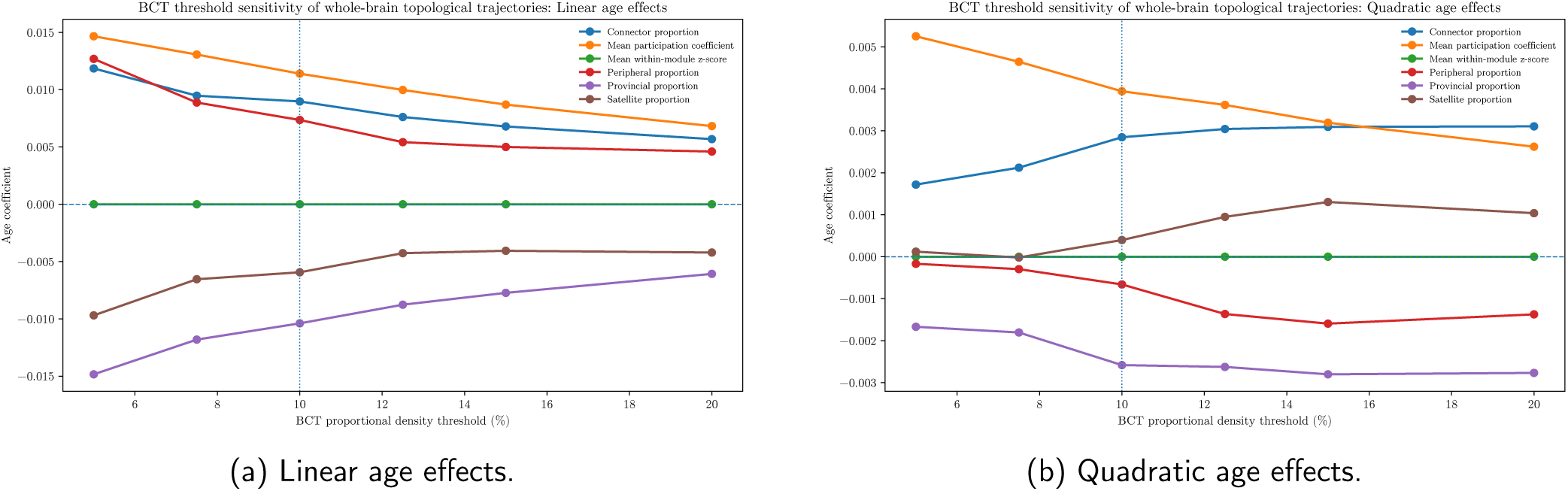
Sensitivity of whole-brain topological age trajectories to the proportional graph-density threshold. *Note.* Analyses were repeated across graph densities from 5% to 20%. The vertical dotted line indicates the primary 10% threshold. The main linear effects were robust across thresholds: connector and peripheral proportions and mean participation coefficient increased with age, whereas provincial and satellite proportions decreased. Quadratic effects were less consistent for the role-proportion models.

The main whole-brain topological findings were not specific to the primary 10% threshold. Across all six thresholds, the linear age effects on connector proportion, peripheral proportion, and mean participation coefficient retained a positive sign and remained FDR-significant. Conversely, provincial and satellite proportions retained negative linear age effects and remained FDR-significant at all thresholds. The whole-brain pattern of increased participation and redistribution away from provincial and satellite roles was therefore robust to threshold choice.

Mean participation coefficient also showed a positive quadratic age component that remained significant across thresholds. By contrast, the quadratic age terms from the binomial role-proportion models were less consistently significant. Network-specific findings were more heterogeneous. Linear age effects on mean participation retained the same direction across thresholds in CON, DMN, FPN, and SMN, but FDR significance varied for some network-specific role proportions. Thus, the broad whole-brain conclusion was robust, whereas finer network-specific role assignments were more sensitive to graph density.

